# TIGIT-Fc Promotes Antitumour Immunity

**DOI:** 10.1101/2020.10.19.346437

**Authors:** Wenyan Fu, Changhai Lei, Jian Zhao, Shi Hu

## Abstract

T cell immunoreceptor with Ig and ITIM domains (TIGIT) is a checkpoint receptor that mediates both T cell and natural killer (NK) cell exhaustion in tumours. An Fc-TIGIT fusion protein was shown to induce an immune-tolerance effect in a previous report, but the relevance of the TIGIT-Fc protein to tumour immunity is unknown. Here, we unexpectedly found that TIGIT-Fc promotes rather than suppresses tumour immunity. TIGIT-Fc treatment promoted the effector function of CD8^+^ T and NK cells in several tumour-bearing mouse models. Additionally, TIGIT-Fc treatment resulted in potent T cell and NK-cell-mediated tumour reactivity, sustained memory-induced immunity in tumour re-challenge models, enhanced therapeutic effects via an antibody against PD-L1, and induction of Th1 development in CD4^+^ T cells. TIGIT-Fc showed a potent antibody-dependent cell-mediated cytotoxicity (ADCC) effect but no intrinsic effect on tumour cell development. Our findings elucidate the unexpected role of TIGIT-Fc in tumour immune reprogramming, suggesting that TIGIT-Fc treatment alone or in combination with other checkpoint receptor blockers is a promising anticancer therapeutic strategy.

## Introduction

T cell immunoreceptor with Ig and ITIM domains (TIGIT, also known as WUCAM, Vstm3 or VSIG9), which is an inhibitory receptor expressed on lymphocytes belonging to the receptor of the Ig superfamily regulatory network that involves multiple players (e.g., CD96/TACTILE, CD112R/PVRIG), is a major emerging target for cancer immunotherapy^1^. As an inhibitor of antitumour responses, TIGIT has the capacity to hinder multiple steps of the cancer immunity cycle, and therefore it is of interest in the development of a first-in-class checkpoint-blocking drug^2^; in addition, new clinical data has shown that combination therapy with the anti-TIGIT antibody tiragolumab combined with atezolizumab appears safe and effective against non-small cell lung cancer (NSCLC)^3^. Several mechanisms of action have been proposed for TIGIT-mediated inhibition of effector T cells and NK cells and the suppression of tumour-specific immunity. TIGIT was reported to interfere with the co-stimulation effect of DNAM-1^4^ or to directly deliver inhibitory signals to the effector cells^5^. Additionally, TIGIT was reported to enhance the suppressive functions of regulatory T cell (Tregs) and hence have the potential to inhibit a wide range of immune cells^6,7^.

TIGIT-Fc is an Fc fusion protein in which the extracellular domain of TIGIT is fused genetically to the immunoglobulin Fc domain. This genetic method enables Fc fusion proteins to have some antibody-like properties, such as long serum half-life and easy expression and purification, making them an attractive platform for research agents and therapeutic drugs^8^. In a previous report, TIGIT-Fc showed an immunosuppressive effect by inhibiting T cell activation in vitro in a dendritic cell-dependent manner and inhibited delayed-type hypersensitivity reactions in mice^9^. We previously also showed that TIGIT-Fc demonstrates a therapeutic effect in a mouse model of lupus^10^. Interestingly, TIGIT-Fc was also shown to promote NK cell activation^11^ and had no effect on cytokine production by tumour-specific CD8^+^ T cells in vitro^12^, indicating that TIGIT-Fc may be a multifaceted immunomodulator and that its function may be dependent on the immune microenvironment. In particular, the impact of TIGIT-Fc on antitumour immunity is currently unknown.

Here, we unexpectedly found that TIGIT-Fc notably reduced the growth of human tumours in a xenograft model containing coimplanted human T cells by supporting a stimulatory immunological mechanism of action for TIGIT-Fc. TIGIT-Fc treatment alone was sufficient to delay tumour growth in vivo and reverse the exhaustion of antitumour T cells and NK cells in multiple tumour models. We further demonstrated that TIGIT-Fc not only directly subverted the exhaustion of tumour-infiltrating NK cells, similar to the effect of blockade of TIGIT with antibodies, but also sustained tumour-specific T cell function in a CD4^+^ T cell-dependent manner. Moreover, a synergistic effect was achieved by the combined therapy of TIGIT-Fc and PD-L1 blockade. TIGIT-Fc also potentially mediated the ADCC lysis of tumour cells in vitro. Hence, these findings demonstrate that TIGIT-Fc may coordinate both the reversal of NK cell exhaustion and the acceleration of the activation of T cells for tumour control and support the clinical development of TIGIT-Fc for cancer immunotherapy.

## Results

### TIGIT-Fc unexpectedly shows antitumour immunity

Targeting co-inhibitory receptors is highly relevant in cancer where therapeutic effects are being exploited clinically. PD-1/PD-L1 blockers are efficacious in the treatment of cancer, but resistance to this therapy is increasing, and the responses to anti-PD-1 or anti-CTLA-4 immunotherapy are relatively minimal in some tumours (such as colorectal carcinomas). Recent reports have shown that anti-TIGIT antibodies have synergistic effects with PD-1 blockade in cancer, both in pre-clinical and clinical studies. As we and others previously have shown that TIGIT-Fc is an immune-suppressive agents^9,10^, we initially aimed to test whether TIGIT-Fc has the capacity to promote resistance to anti-PD-L1 therapy in xenograft models of the human melanoma cell line A375. Both TIGIT-Fc and atezolizumab were confirmed to bind to A375 cells (Fig. 1a). To generate allogenic T cell lines with specificity to A375 cells^13^, primary human T cells were expanded in culture and implanted subcutaneously in NSG mice together with tumour cells. Our data shows that the administration of the anti-PD-L1 antibody atezolizumab led to a notable growth inhibition of the A375 tumour in the presence of human T cells, consistent with a previous report^13^. Unexpectedly, TIGIT-Fc alone significantly inhibited the tumour growth of A375 xenografts compared with an isotype-matched control antibody. Moreover, the combined therapy of TIGIT-Fc and atezolizumab therapy showed a superior tumour growth inhibition reaching nearly 100% (Fig. 1b). Based on the immunoglobulin-like structure of TIGIT-Fc, we hypothesize that TIGIT-Fc may have a two-sided biological effect in that it is a potent agonist of CD155 and a membrane-bound TIGIT antagonist. We tested the antitumour effect of the anti-TIGIT antibody etigilimab in the T cell-based A375 tumour model, but etigilimab alone showed no inhibitory effect on the growth of the A375 tumour xenograft, and combined therapy with etigilimab and atezolizumab showed no synergistic effect compared with atezolizumab monotherapy (Fig. 1c), suggesting that the antitumour efficacy of TIGIT-Fc is not due to TIGIT blockade in A375 xenografts.

**Figure 1.**
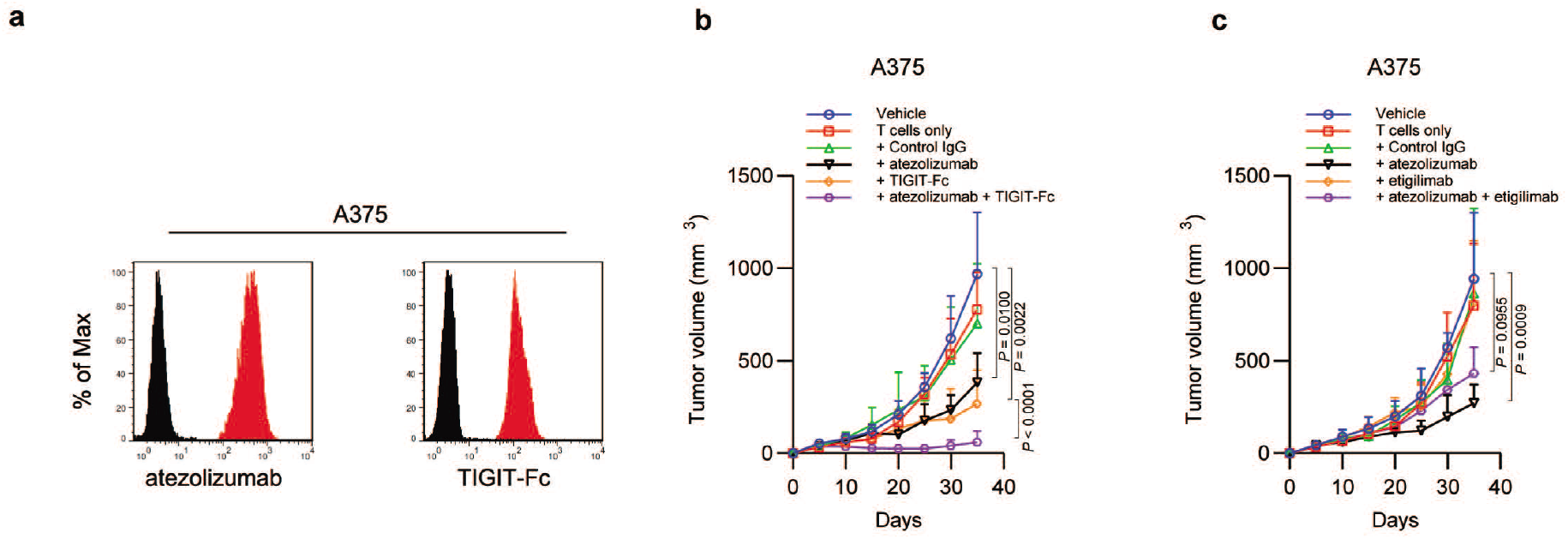
TIGIT-Fc shows antitumour activity in xenograft mouse models of human cancer. **a.** The binding of atezolizumab and TIGIT-Fc to A375 cells was detected by flow cytometry analysis. The histograms shown in black correspond to the isotype controls, whereas the red histograms indicate the positive fluorescence. **b-c.** Tumour volumes of A375 in NOD/SCID mice following coimplantation of primary human T cells for assessing the therapeutic effect of atezolizumab in combination with TIGIT-Fc (b) or an anti-TIGIT antibody (c). Antibodies were used in a dose of 10 mg/kg alone or in combination (5 mg/kg each), as specified in the figure. The data are the means ± s.d., *n* = 8. *P* values were generated by two-way ANOVA followed by a Bonferroni posttest comparison.

### TIGIT-Fc enhances effector NK cell function in tumour-bearing mice

A previous report showed that blockade of TIGIT prevents NK cell exhaustion^14^. To assess whether TIGIT-Fc can blockade TIGIT on NK cells similar to that of an anti-TIGIT antibody in tumour-bearing mice, murine TIGIT-Fc (mTIGIT-Fc) was first tested against subcutaneous MC38 colon adenocarcinoma tumours, since this model is considered a standard and has been demonstrated by many laboratories as anti-PDL1-sensitive and immunogenic.

In the MC38 tumour model (Fig. 2a), we found that tumour growth was notably inhibited by the administration of mTIGIT-Fc, as shown by the lower tumour volume (Fig. 2a) and the improved overall survival of mice (Fig. 2b). Moreover, three of eight mice with the mTIGIT-Fc treatment showed complete tumour regression, suggesting that TIGIT-Fc has the capacity to induce a strong antitumour immunity. Administration of mTIGIT-Fc resulted in a higher frequency of CD107a^+^, TNF^+^, and IFN-γ^+^ tumour-infiltrating NK cells than in mice treated with the control IgG (Fig. 2c), indicating that mTIGIT-Fc reversed the exhaustion of the tumour-infiltrating NK cells. Of note, a significantly higher frequency of tumour-infiltrating CD8^+^ T cells with surface expression of CD107a, TNF and IFN-γ was also observed to accompany the mTIGIT-Fc treatment. Another colorectal carcinoma mouse model, CT26, was also employed in our study, in which it was observed that mTIGIT-Fc delayed tumour growth and prolonged overall mouse survival (Fig. 2e-f), and two of eight mice showed complete tumour regression with the mTIGIT-Fc treatment. Consistent with the observation in MC38 tumours, mTIGIT-Fc alleviated NK cell exhaustion, as a higher frequency of tumour-infiltrating NK cells expressing CD107a, TNF and IFN-γ were observed. Additionally, mTIGIT-Fc treatment showed a higher frequency of tumour-infiltrating CD8^+^ T cells expressing TNF and IFN-γ. Thus, these results indicated that TIGIT-Fc shows a similar TIGIT blockade effect as an anti-TIGIT antibody to prevent the exhaustion of NK cells in tumour-bearing mice with a beneficial effect on T cells.

**Figure 2.**
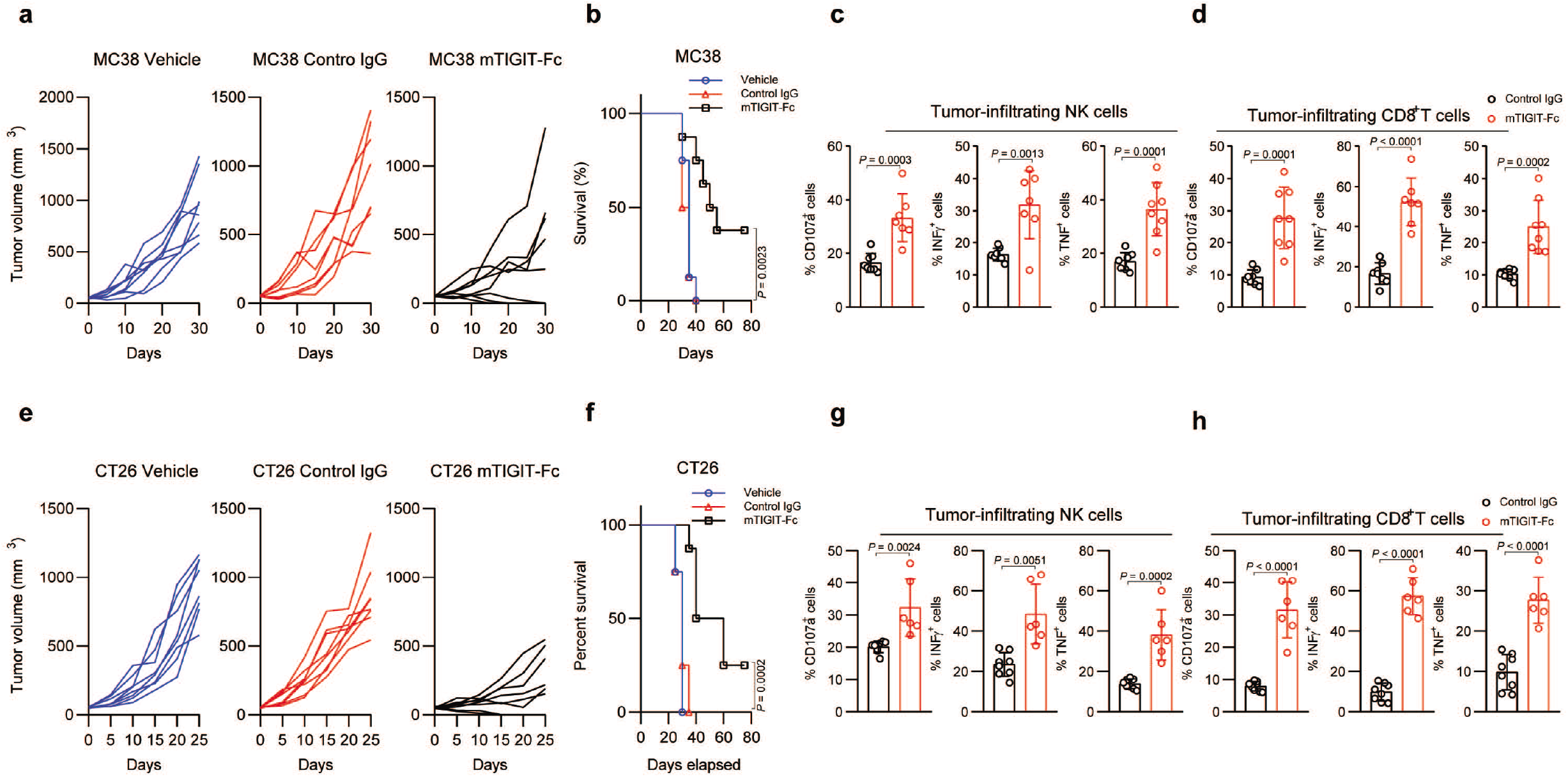
TIGIT-Fc inhibits tumour growth and prevents exhaustion of tumour-infiltrating NK cells. **a.** Tumour volumes of different MC38 tumour xenografts after the indicated weekly treatment with mTIGIT-Fc (10 mg/kg) or control IgG, as specified in the figure. **b**. Survival of mice as in **a** at various times. *P* values, Mantel-Cox test. **c-d**. Frequency of cells expressing CD107a, TNF, IFN-γ or CD226 among tumour-infiltrating NK cells (**c**) or T cells (**d**) in mice as in **a** (n = 6-8 per group), assessed when the tumours in the mice treated with mTIGIT-Fc reached a size of ~250 mm^3^. **e**. Tumour volumes of different CT26 tumour xenografts after the indicated weekly treatment with mTIGIT-Fc (10 mg/kg) or control IgG, as specified in the figure. **f**. Survival of mice as in **e** at various times. *P* values, Mantel-Cox test. **g-h**. Frequency of cells expressing CD107a, TNF, IFN-γ or CD226 among tumour-infiltrating NK cells (**g**) or T cells (**h**) in mice as in **e** (n = 6-8 per group).

### TIGIT-Fc has a strong antitumour effect on CD155-positive tumours in the absence of adaptive immunity

We then determined whether the antitumour effect of TIGIT-Fc can occur in nude mice that lack host T cells. In CT26 tumour model based on nude mice, we found that following treatment with mTIGIT-Fc, CT26 tumour growth was notably inhibited (Fig. 3a), and the tumour-infiltrating NK cells showed improved function, as indicated by the increased frequency of such cells expressing CD107a, TNF or IFN-γ in the nude mice model (Fig. 3b). The therapeutic value of the recombinant human TIGIT-Fc-targeting regimen was further assessed in mice xenografted with human cancer cells. As shown in Fig. 4c, the vehicle A431 tumours progressed rapidly and reached volumes of greater than 1,000 mm^3^ in less than 30 days. Conversely, TIGIT-Fc treatment effectively delayed tumour growth to this volume for approximately 40 days. TIGIT-Fc treatments also efficiently inhibited tumour growth and delayed tumour growth for 45 days and 30 days in SK-BR-3 or SK-OV-3 tumours, respectively (Fig. 3c). As we detected high CD155 expression in the mouse tumour cell line CT26 and in the human tumour cell line A431, SK-BR-3 cells and SK-OV-3 cells (Fig. 4d), we therefore use H2126 cells with very low detectable CD155 expression to further evaluate the antitumour role of TIGIT-Fc.

**Figure 3.**
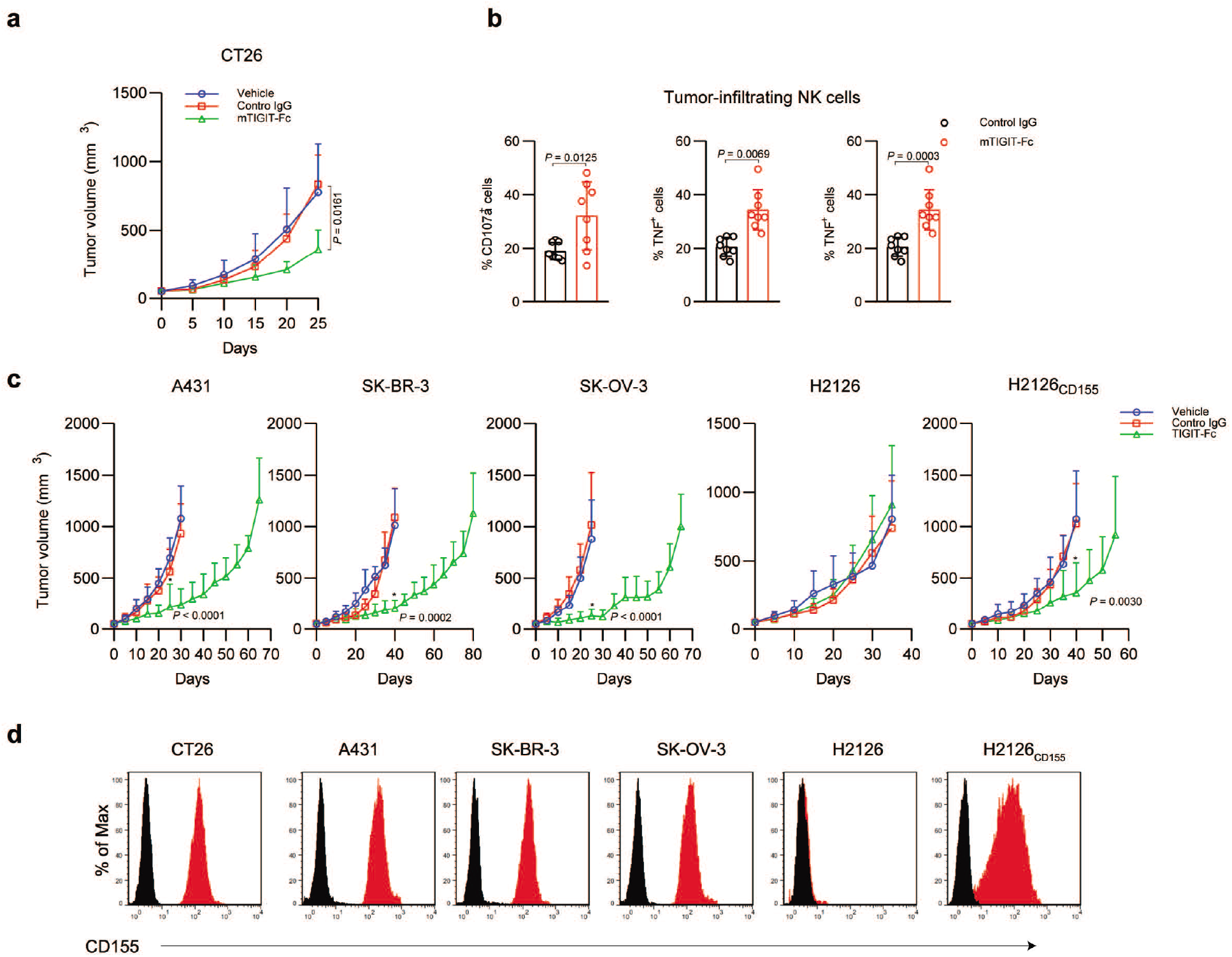
TIGIT has a protective role in mice with adaptive immunodeficiency. **a** Tumour volumes of different CT26 tumour xenografts in nude mice after the indicated weekly treatment with mTIGIT-Fc (10 mg/kg) or control IgG, as specified in the figure. **b**. Frequency of cells expressing CD107a, TNF, or IFN-γ among tumour-infiltrating NK cells in mice as in **a** (n = 6-8 per group), assessed when tumours in mice treated with mTIGIT-Fc reached a size of ~250 mm^3^. **c**. Tumour volumes of different A431, SK-BR-3, SK-OV-3, H2126 or H2126-CD155 tumour xenografts in nude mice after the indicated weekly treatment with TIGIT-Fc (10 mg/kg) or control IgG, as specified in the figure. **d**. The expression of CD155 in different tumour cell lines was detected by staining with an anti-CD155 antibody, followed by flow cytometry analysis. The histograms shown in black correspond to the isotype controls, whereas the red histograms indicate the positive fluorescence.

Interestingly, we observed that the therapeutic benefit of TIGIT-Fc treatment on tumour growth in the CD155 low-expression model was minimal (Fig. 4c). The CD155 expression of tumour cells contributing to the antitumour effect of TIGIT-Fc was further supported in the tumour model of H2126 cell derivatives engineered to express CD155, in which TIGIT-Fc showed a notable antitumour effect in tumour-bearing mice. In all the tumour models based on nude mice, despite the delayed tumour growth and survival advantage, mice treated with TIGIT-Fc eventually succumbed to the tumours, and no mice had the capacity to reject tumour cells, suggesting that the antitumour effect of TIGIT-Fc would be weakened by dysfunction of the host T cells.

**Figure 4.**
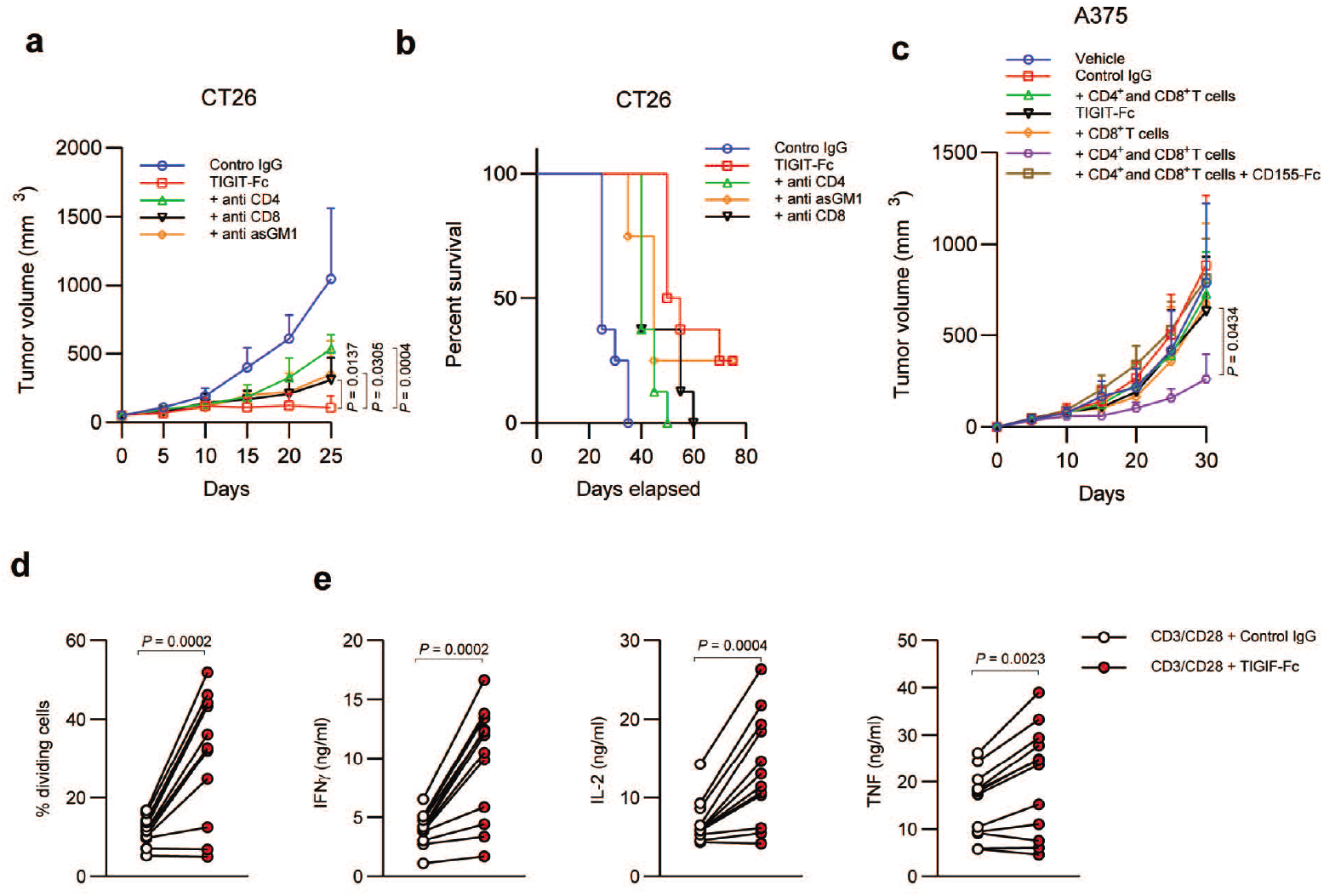
CD4+ T cells contribute to the antitumour efficacy of TIGIT-Fc. a. Tumour volumes of different CT26 tumour xenografts after the indicated weekly treatment with mTIGIT-Fc (10 mg/kg) or control IgG, or plus depletion of different immune cells as specified in the figure: anti-CD8 (53.5.8, CD8^+^ T cell depletion); anti-CD4 (GK1.5, CD4^+^ T cell depletion), anti-asGM1 (NK cell depletion) (antibodies were given weekly at the dose of 10 mg/kg). **b**. Survival of mice in **a. c**. Tumour volumes of A375 in NSG mice following coimplantation of different primary human T cells for assessing the therapeutic effect of TIGIT-Fc. **d-e**. proliferation assay by CFSE dilution (**d**) and ELISAs for IL-2, IFN-γ, and TNF production (**e**) after stimulation of PMBCs with plate-bound anti-CD3/CD28 plus TIGIT-Fc or control IgG.

### CD4^+^ T cells have a pivotal role in the antitumour immunity of TIGIT-Fc

To further explore the mechanism that underlies the antitumour immunity of TIGIT-Fc, mice were depleted of NK cells, CD4^+^ T cells, and CD8^+^ T cells in combination with mTIGIT-Fc treatment in syngeneic immune-competent mouse models. In CT26 tumour models, we found that the depletion of CD4^+^ T cells, CD8^+^ T cells or NK cells significant inhibited the therapeutic effect of mTIGIT-Fc (Fig. 4a-b). A previous report showed that depletion of NK cells abolished the therapeutic effect of anti-TIGIT antibodies, and interestingly, two of eight mice rejected tumours after treatment with TIGIT-Fc combined with an anti-asialoGM1 (anti-asGM1) antibody, suggesting that TIGIT-Fc has NK-independent antitumour mechanisms.

Notably, the tumour growth effect of TIGIT-Fc, which was significantly weakened in the CD4^+^ T cell deficiency context, indicated a unique role of CD4^+^ cells in the setting of TIGIT-Fc treatment. To further assess the impact of T cells on the therapeutic effect of TIGIT-Fc, CD8^+^ cells or CD4^+^ and CD8^+^ T cells were provided by coimplanting human cells into NSG mice bearing A375 or tumours along with different treatments. Our results demonstrated that reduced tumour growth was only observed in mice treated with TIGIT-Fc together with both CD4^+^ and CD8^+^ T cells, whereas the impact of the TIGIT-Fc therapy with CD8^+^ T cells alone was minimal (Fig. 4c). Moreover, additional treatment with recombinant CD155 abolished the antitumour effect of TIGIT-Fc, suggesting that the T cell-based antitumour effect of TIGIT-Fc is dependent on CD155. Notably, functional assessment of TIGIT-Fc in vitro in anti-CD3/anti-CD28-stimulated PBMCs from various donors demonstrated the potent ability of this fusion protein to enhance CD4^+^ and CD8^+^ T cell proliferation (Fig. 4d). TIGIT-Fc also significantly increased Th1 cytokine production (IFN-γ, TNF-α and IL-2) in these cultures compared with a control IgG (Fig. 4e).

### TIGIT-Fc elicits a potent anti-tumour memory response and enhances PD-L1 blockade

Our data above showed that TIGIT-Fc has the capacity to evoke potent antitumour immunity involving both T cells and NK cells. We then tested whether the mice that survived with TIGIT-Fc treatment after tumour challenge have anamnestic responses to tumour re-challenge. MC38 tumour-bearing mice that were initially treated with mTIGIT-Fc and experienced a complete response were subsequently given a second implantation of MC38 cells in the absence of any further treatment (Fig. 5a). Notably, the MC38 rechallenge was incapable of establishing a tumour mass in mice previously cured of tumours, while tumour-naive mice with no previous treatment showed a successful tumour engrafting. Similar results were obtained in CT26 tumour-bearing mice (Fig. 5b) cured with mTIGIT-Fc treatment, where a rechallenge with tumour cells showed no sign of tumour growth in the mice. These results demonstrated that TIGIT-Fc mono-immunotherapy has the capacity to elicit a strong antitumour immune memory.

**Figure 5.**
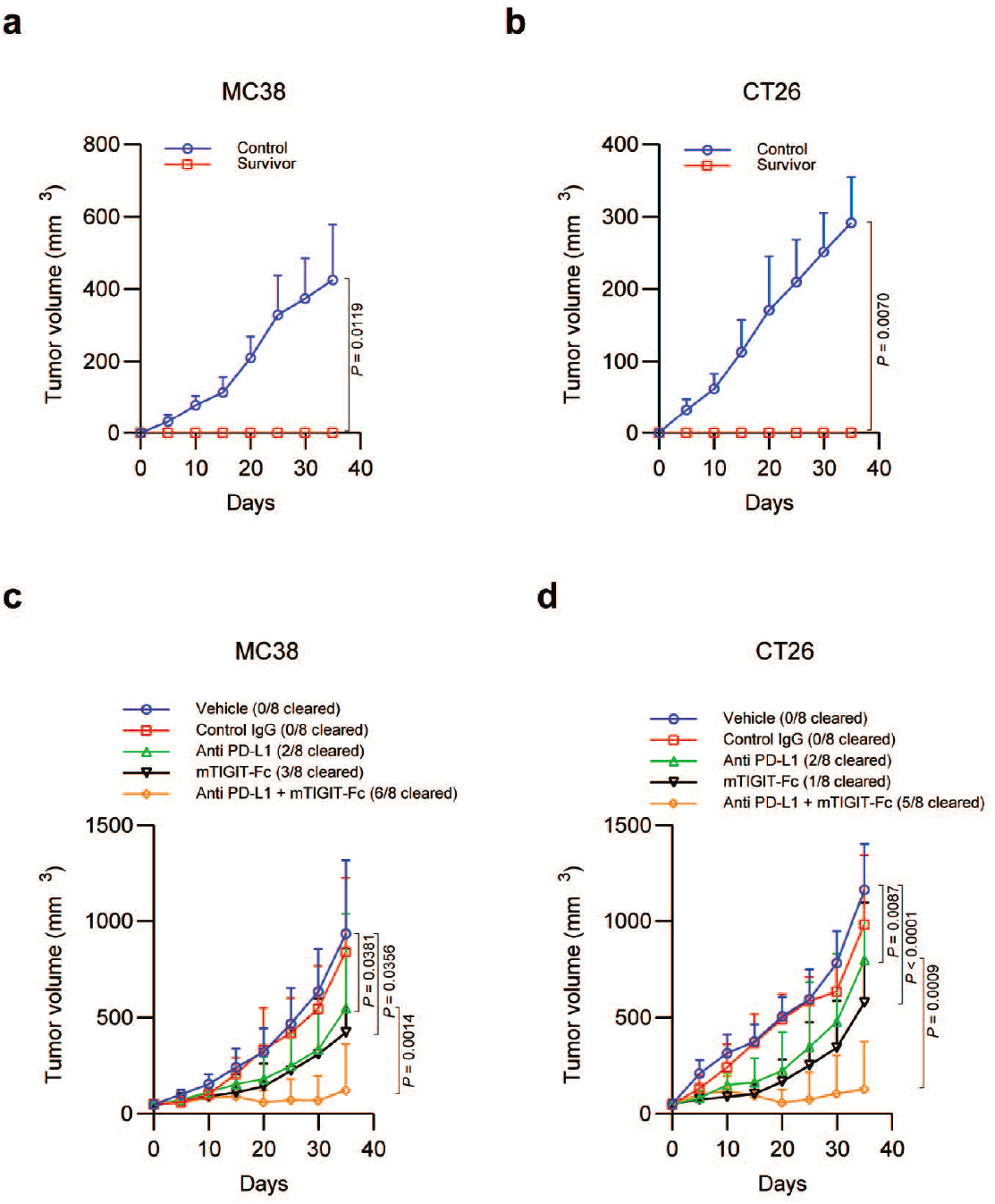
TIGIT-Fc induces an antitumour memory response and shows a synergic effect with anti-PD-L1. **a-b.** Tumour rechallenge experiments. The surviving mice xenografted with MC38 (**a**) or CT26 (**b**) after mTIGIT-Fc treatment were rechallenged with different tumours. **c-d**. Tumour volumes of different MC38 (**c**) or CT26 (**d**) tumour xenografts after the indicated weekly treatment with mTIGIT-Fc (10 mg/kg) or control IgG, as specified in the figure.

The combination of TIGIT-Fc and atezolizumab that showed a strong antitumour effect in the human T cell-based A375 model is of interest. We therefore investigated whether the combination of mTIGIT-Fc and anti-PD-L1 antibody would enhance the therapeutic outcome by assessing the potential efficacy of the combined therapy in mice with established syngeneic tumours. In both MC38 and CT26 colorectal adenocarcinoma (Fig. 5c-d), treatment with mTIGIT-Fc or anti-PD-L1 monotherapy resulted in reduced tumour growth, with two of eight and three of eight mice tumour-free after 35 days (25%–37.5%), respectively. Following combinatorial TIGIT-Fc/anti-PD-L1 immunotherapy, 75% and 62.5% of the MC38- and CT26-inoculated mice, respectively, were tumour-free after 35 days Thus, these data showed that the combined therapy of TIGIT-Fc and anti-PD-L1 induces potent antitumour effects, leading to improved tumour control.

### TIGIT-Fc has a potent ADCC effect but no intrinsic effect on tumour cells

TIGIT-Fc itself may have potent ADCC effect because it is an Fc-fused protein, and therefore we evaluated the ability of TIGIT-Fc to induce ADCC lysis of human tumour cell targets expressing CD155 in vitro utilizing PBMC-dependent ADCC assays. Our data shows that the CD155 high-expressing cell lines A431, SK-BR-3 and SK-OV-3 were sensitive to ADCC-mediated lysis by TIGIT-Fc (Fig. 6a) over a range of E: T ratios, whereas the CD155-negative cell line H2126 was resistant to lysis. We next investigated whether TIGIT-Fc has an intrinsic effect on tumour cells. TIGIT-Fc treatment was evaluated for effects on in vitro cell viability, migration, and invasion in the CD155 high-expressing cells. However, no effect on the in vitro proliferation of both SK-BR-3 and SK-OV-3 cells was observed after TIGIT-Fc treatment compared with control IgG (Fig. 6b). In migration, invasion and colony formation assays, TIGIT-Fc treatment also showed a negligible effect on tumour cells (Fig. 6c-d). To confirm these findings, the effects of TIGIT-Fc on the in vivo tumour growth of tumour cells were further examined. Tumour growth was monitored in subcutaneous implantation NSG mice models to exclude the impact of immune cells. Our data showed that in the NSG mice, TIGIT-Fc treatment had no detectable effect of tumour growth in both A431 cells and SK-BR-3 cells (Fig. 6e).

**Figure 6.**
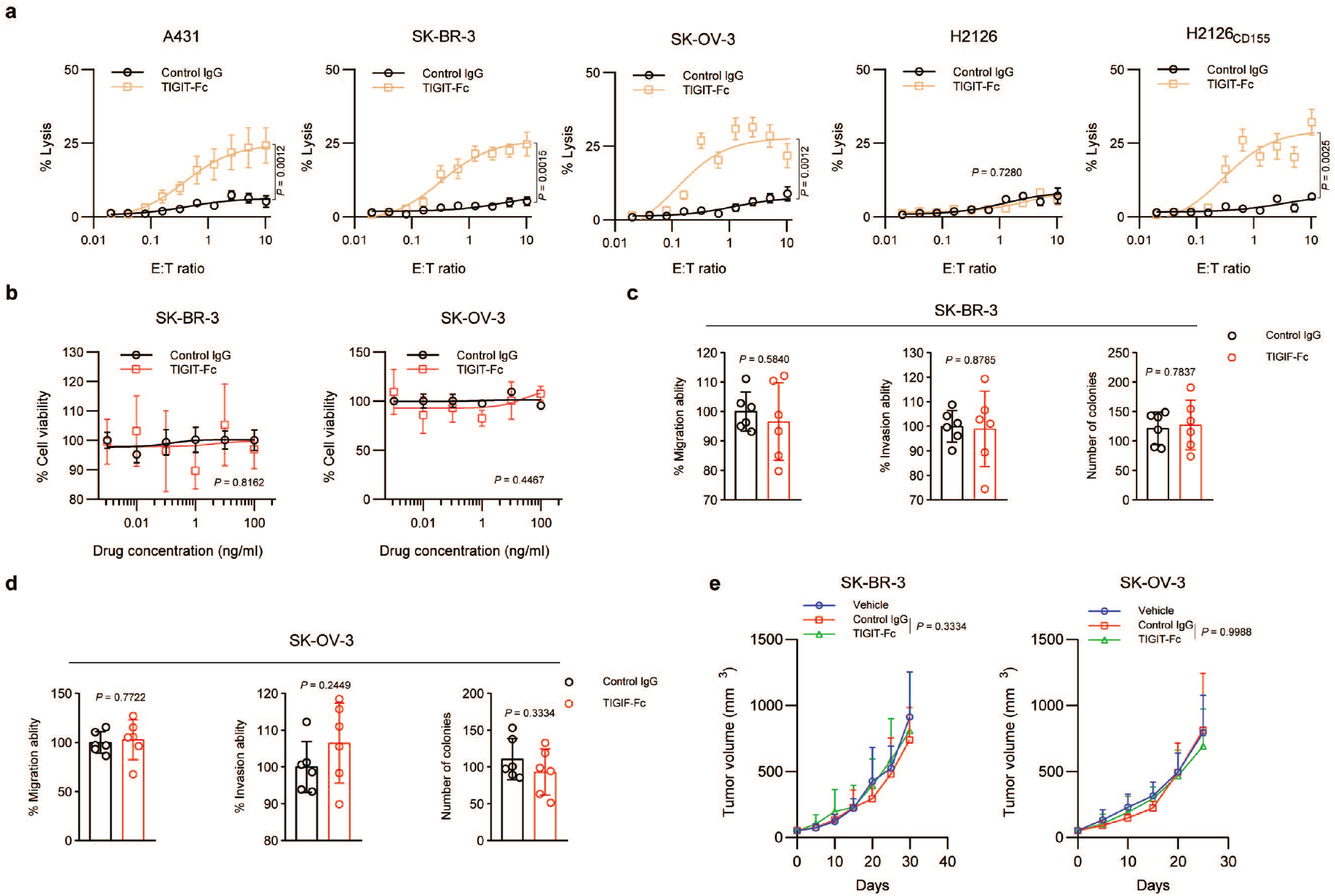
TIGIT-Fc shows an ADCC effect but no intrinsic effect on tumour cells. **A.** In vitro ADCC assay using tumour cell lines with different levels of CD155 expression as targets. **b.** The effects of TIGIT-Fc on the in vitro viability of different tumour cells. **c-d**. The effects of TIGIT-Fc on migration, invasion and colony formation of SK-BR-3 (**c**) and SK-OV-3 cells (**d**). **e**. Tumour volumes of different SK-BR-3 or SK-OV-3 tumour xenografts in NSG mice after the indicated treatment.

## Discussion

TIGIT-Fc was used to simulate membrane-bound TIGIT function in an earlier study^9^. Upon binding with TIGIT-Fc, the activation of CD155 in human monocyte-derived DCs (MDDCs) led to decreased secretion of the proinflammatory cytokine IL-12 and increased secretion of IL-10. It should be noted that in addition to TIGIT-Fc, CD226-Fc also has been shown to modify DC cytokine production, suggesting that this effect is mainly caused by CD155 signalling in MDDCs^9^. Interestingly, subsequent studies reported that agonistic anti-TIGIT mAbs inhibit anti-CD3/anti-CD28 mAb-mediated T cell proliferation and cytokine production in the absence of APCs in humans and mice^15–17^. A recent study further found that the IFN-γ production of CD8+ T cells was suppressed by melanoma cells expressing a truncated version of CD155 in a similar manner as cells expressing wild-type CD155^18^. Taken together, these studies suggested that the TIGIT-CD155 interaction can inhibit T cell functions without downstream signalling via CD155, highlighting the cell-intrinsic mechanisms of TIGIT. The observation that inhibitory signals can also be directly transmitted via the cytoplasmic tail of TIGIT further supported its cell-intrinsic role. Two publications from the same group established that the ITIM motif is essential for human TIGIT signalling, whereas mouse TIGIT inhibition can be mediated by either the ITIM motif or the ITT motif alone^5,19^. Moreover, another group suggested an important role for the intracellular ITT-like motif in human TIGIT and highlighted two different signalling pathways interfering with NK cell cytotoxicity or IFN-γ production^20,21^. Additionally, whole-genome microarray analysis showed that mouse T cell activation was suppressed by TIGIT engagement by downregulating T cell receptor (TCR) expression, together with several other molecules involved in TCR and CD28 signalling^17^.

Structurally, TIGIT-Fc is an immunoadhesin, which is an immunoglobulin (Ig)-like chimeric protein comprised of target-binding regions fused to the Fc-hinge region of Ig and is designed to have a long half-life and antibody-like properties. Mechanistically, TIGIT-Fc has the capacity to block TIGIT-associated interactions, including the binding of CD155-TIGIT. In a xenograft model system to investigate the T cell-mediated tumour cell killing effect, administration of TIGIT-Fc in A375 xenografts resulted notably in tumour growth inhibition. Importantly, the antitumour effect of TIGIT-Fc was entirely dependent upon the presence of the tumour-reactive human T cells in this model. These data unexpectedly demonstrated that the ability of TIGIT-Fc increased tumour cell elimination by T cells and supported the further investigation of TIGIT-Fc for the treatment of cancer. TIGIT-Fc treatment enhanced NK cell-mediated antitumour immunity both in syngeneic tumour models and tumour xenograft models in nude mice, which is very similar to the effect of an anti-TIGIT antibody^14^, demonstrating that the blockade of TIGIT signalling of tumour-infiltrating NK cells by TIGIT-Fc can be achieved. Interestingly, unlike the anti-TIGIT antibody, which promotes tumour-specific T cell immunity in an NK cell-dependent manner, TIGIT-Fc exhibited a direct activation effect on tumour-specific T cells. We further found that CD4^+^ T cells were critical for the therapeutic effects of TIGIT-Fc and that TIGIT-Fc treatment augmented proliferation of T cells and mediated Th1 development from naive CD4^+^ T cells in vitro. These data are consistent with a previous study that showed that agonist anti-CD155 antibodies induced Th1 development in both humans and mice, as evidenced by production of IFN-γ and upregulation of *Tbx21* transcription^22^. It should be noted that T cells derived from different healthy donor show different responses to TIGIT-Fc (Fig. 4d-e), indicating a heterogeneity in CD155 signalling-based T cell function.

CD155 is a member of the poliovirus receptor-related (PRR) family of adhesion molecules, along with CD111 (nectin-1/PRR-1), CD112 (nectin-2/PRR-2), nectin-3 (PRR-3), and nectin-4 (PRR-4). CD155 was initially identified as a receptor for poliovirus in humans^23^ and binds to another PRR family member, nectin-3, as well as to the matrix protein vitronectin, thereby mediating cell-cell or cell-matrix adhesion, respectively, and cell migration^24^. Additionally, CD155 is a ligand for CD226, CD96, and TIGIT on T cells and NK cells. The expression of CD155 in normal tissue, such as epithelial and endothelial cells, is very low, but most tumour cells express high levels of CD155 ^24^. Dysregulation of CD155 expression promotes tumour cell invasion, migration, and proliferation and is associated with a poor prognosis and enhanced tumour progression^24–26^, but the definition of tumour cell-intrinsic and -extrinsic effects of CD155 has not been fully explored. In present study, we show that targeting CD155 with TIGIT-Fc did not show an effect on the development of tumour cells in vitro, indicating that at least CD155 did not directly participate in the oncogenic signalling in tumour cells. Moreover, TIGIT-Fc showed a potent ADCC effect, which is well established as a major mode of tumour-cell killing with immune cells, paving the way for TIGIT-Fc as a possible therapeutic candidate for CD155-overexpressing tumours.

Our study has several limitations. We provide evidence that promoting tumour-specific immunity with TIGIT-Fc can be achieved, but our in vivo efficacy models may not fully recapitulate human tumours, and the data are from a small number of animals. Moreover, the mechanisms responsible for these therapeutic effects of TIGIT-Fc are currently not well characterized. As some other cells in the tumour microenvironment also express CD155, e.g., tumour-infiltrating myeloid cells, the mechanisms of TIGIT-Fc may be more complicated. Hence, these findings will need further validation.

To sum up, our study reveals that TIGIT-Fc can induce the antitumour immune response of NK cells and T cells. Based on the wide application of T cell- and NK cell-based treatment strategies in clinical practice, targeting agents that intervene in the CD155-TIGIT axis hold translational potential. Our results supported the conclusion that treatment with TIGIT-Fc alone or in combination with other immune checkpoint inhibitors is a promising strategy to improve the outcomes of tumour immunotherapy.

## Methods

### Cell lines, primary cells, and therapeutic IgGs

The human cell lines A375, A431, SK-BR-3, SK-OV-3, and H2126 were purchased from the American Type Culture Collection (ATCC, Manassas, VA). The identities of the cell lines were verified by STR analysis. CT26 cells were purchased from the Chinese Academy of Sciences (Shanghai, China). MC38 cells purchased from Kerafast, Inc. All cell lines were confirmed to be mycoplasma-free. The cells were maintained in DMEM with 10% foetal bovine serum. Human peripheral blood mononuclear cells (PBMC) were isolated from healthy volunteer donors by layering over Ficoll-Paque Plus (GE Healthcare) as per the manufacturer’s instructions and were enriched for CD4^+^ or CD8^+^ T cells using RosetteSep (STEMCELL Technologies) T cell enrichment as per the manufacturer’s instructions. All specimens were collected under an approved protocol by the Second Military Medical University Review Board, and written informed consent was obtained from each donor. The fusion proteins used were as previously described^10^; briefly, a recombinant plasmid was constructed by fusing the Fc segment of human IgG1 or murine IG2a, encoding the hinge-CH2-CH3 segment, to the C-termini of the extracellular domains (ECDs) of human and murine TIGIT, respectively. All fusion proteins were obtained via the FreeStyle 293 Expression System (Invitrogen) and subsequently purified using protein A-sepharose from the harvested cell culture supernatant. The purity of the fusion protein was determined by polyacrylamide gel electrophoresis. The protein concentration was measured according to the UV absorbance at a wavelength of 280 nm.

### In vivo mice study

Mice were housed in a specific pathogen-free barrier facility. All the in vivo experiments were approved by the Institutional Animal Care and Use Committee (IACUC) of Second Military Medical University. For mouse models with human T cells, tumour-reactive T cells were expanded in culture with mitomycin C-treated cancer cells for 10 days in RPMI 1640 supplemented with 10% FBS and IL2. Tumour-reactive T cells or subsets were mixed with A375 cells at a 1:5 ratio, and the cell mixtures were implanted by subcutaneous injection of 5 ×10^6^ cells into the flank of NSG mice (Shanghai Model Organisms Center, Inc.). Different drugs were administered i.p. 1 hr immediately after implantation. For syngeneic tumour models, MC38 or CT26 cells were inoculated into C57BL/6 mice or BALB/c mice. For tumour xenograft models, CT26, A431, SK-BR-3 or SK-OV-3 cells were inoculated into BALB/c nude mice (Shanghai Experimental Animal Center of Chinese Academy of Sciences) or NSG mice. When tumour volumes reached an average of approximately 50 mm^3^, the mice were randomly divided into groups of 8 mice each. Multiple dose studies consisted of 3 weeks of treatment, with a 2× loading dose on the first dose (day of randomization). Tumour volumes were calculated by the following formula: volume = length × (width)^2^/2.

### Isolation of tumour-infiltrating lymphocytes (TILs)

Tumour tissues were dissociated, and TILs were isolated in the presence of collagenase I (0.1% w/v, Sigma) and DNAse (0.005% w/v, Sigma) for 1 h before centrifugation on a discontinuous Percoll gradient (GE Healthcare). Isolated cells were then used in various cell function assays

### Flow cytometry

Cell surface staining was performed at 4 °C for 30 min and was analysed using a FACSCalibur flow cytometer (BD Biosciences) and CellQuest Software (BD Biosciences). A minimum of 1 × 10^4^ cells was examined. Antibodies to mouse CD3ε (145-2C11, BD Bioscience), CD107a (1D4B, BD Bioscience), IFN-γ (XMG1.2, BioLegend), TNF (MP6-XT22, BioLegend), CD155 (4.24.1, BioLegend), CD8α (53-6.7, BioLegend) and antibodies to human CD3 (HIT3a, BioLegend), CD45 (2D1, BioLegend), CD4 (RPA-T4, BD Bioscience), and CD8 (RPA-T8, BD Bioscience) were used in this study. For TILs experiments, the cells were stimulated with 30 ng/ml phorbol 12-myristate 13-acetate (PMA, Sigma) and 1 μM ionomycin (Sigma) in the presence of 2.5 μg/ml monensin (eBioscience) for 4 h, after which they were stained for surface markers and fixed and permeabilized with eBioscience FoxP3 fixation buffer according to the manufacturer’s instructions. Fixed cells were stained with antibodies to IFN-γ (XMG1.2; BioLegend) and TNF (MP6-XT22, BioLegend).

### In vitro ADCC assay

Target cells were labelled with ^51^Cr for 1 h at 37 °C. Radioactive ^51^Cr (50 μCi) was used to label 1 × 10^6^ target cells. Labelled target cells (n = 5000) were plated in each well of a 96-well plate in a volume of one hundred microliters. Effector cells were added at a volume of 100 μl at different E:T ratios with different drugs. The cells were incubated together for 4 h at 37 °C. The supernatant (30 μl) from each well was collected and transferred to the filter of a LumaPlate. The filter was allowed to dry overnight. Radioactivity released into the culture medium was measured using a β-emission-reading liquid scintillation counter. The percentage of specific lysis was calculated as follows: (sample counts – spontaneous counts)/(maximum counts – spontaneous counts) × 100.

### In vitro tumour cell line assay

For cell viability assay, tumour cells were plated in triplicate at 5× 10^3^ per well in 96-well plates overnight. After plating, the cell culture medium was replaced with assay medium containing 0.1% FBS. TIGIT-Fc and control IgG were added at multiple concentrations, and 72 hours later, cell viability was measured using the CellTiter-Glo Luminescent Cell Viability Assay (Promega). For the migration assay, cell mobility was assessed with a modified two-chamber migration assay (8-mm pore size, BD Biosciences, Bedford, MA) according to the manufacturer’s instructions as described previously^27^ Briefly, approximately 2 × 10^4^ cells were plated into the upper chamber for 18–24 hours along with different treatments (1 mg/ml) and allowed to migrate into the lower chamber. The cells at the bottom of the membrane were fixed and stained with 20% methanol/0.2% crystal violet, while the cells in the upper chamber were removed using cotton swabs. The invasion assay was performed as previously described using transwell cell culture chambers (8-μm pore size polycarbonate membrane, Costar) ^28^. Briefly, 100 μl of cell suspension at 1 × 10^6^ cells/ml with different treatments (1 mg/ml) in DMEM supplemented with 0.5% foetal bovine serum was loaded into the upper chamber, and the lower chamber was loaded with 600 μl of DMEM with 10% foetal bovine serum. The membranes were precoated with Matrigel (BD Pharmingen). The chamber plates were incubated at 37°C for 24 hours. Then, the filter was fixed in 4% paraformaldehyde and stained with haematoxylin (Sigma). The cells on the upper side of the filter were removed with a cotton swab, and the cells that had passed through the membrane were counted in 10 randomly selected microscopic fields. For the colony formation assay, the cells were seeded onto 6-well plates or 3.5-cm dishes along with different treatments (1 mg/ml). Colonies were allowed to form in an incubator at 37°C and 5% CO_2_ for 10 days. At the end of the incubation period, the clones were fixed and stained with 0.5% crystal violet, and colonies larger than 50 μm in diameter were counted.

### Statistical analysis

Unless otherwise specified, the Student’s t-test was used to evaluate the significance of differences between 2 groups, and ANOVA was used to evaluate differences among 3 or more groups. Differences between samples were considered significant when *P* < 0.05.

## Acknowledgements

This study was supported by the National Natural Science Foundation of China (grant nos. 81773261, 31970882, 81903140 and 81602690); the Shanghai Rising-Star Program (grant no. 19QA1411400); the Shanghai Sailing Program (19YF1438600); and the Shanghai Chenguang Program (grant no. 17CG35).

